# Boosting learning efficacy with non-invasive brain stimulation in intact and brain-damaged humans

**DOI:** 10.1101/500215

**Authors:** F. Herpich, M.D. Melnick, S. Agosta, K.R. Huxlin, D. Tadin, L. Battelli

## Abstract

Numerous behavioral studies have shown that visual function can improve with training, although perceptual refinements generally require weeks to months of training to attain. This, along with questions about long-term retention of learning, limits practical and clinical applications of many such paradigms. Here, we show for the first time that *just 10 days* of visual training coupled with transcranial random noise stimulation (tRNS) over visual areas causes dramatic improvements in visual motion perception. Relative to control conditions and anodal stimulation, tRNS-enhanced learning was at least twice as fast, and, crucially, it persisted for 6 months after the end of training and stimulation. Notably, tRNS also boosted learning in patients with chronic cortical blindness, leading to recovery of motion processing in the blind field after just 10 days of training, a period too short to elicit enhancements with training alone. In sum, our results reveal a remarkable enhancement of the capacity for long-lasting plastic and restorative changes when a neuromodulatory intervention is coupled with visual training.

## Introduction

The human brain changes throughout life (Gilbert & Li, 2012; Liat al., 2004). Visual training is a well-known tool for inducing such changes, improving sensory performance in healthy adults (reviewed in Dosher & Lu, 2017; Li, 2016; Sagi, 2011; Wang et al., 2016); and in various clinical populations (Deveau et al., 2013; Melnick et al., 2016; Nyquist et al., 2016), a phenomenon referred to as visual perceptual learning (VPL). The specific role of different visual areas of the brain during VPL is still openly debated, with several mechanisms likely contributing to learning. For instance, neurophysiological studies have shown that perceptual learning selectively modifies the signal strength of neurons responding to relevant stimulus features, while concurrently suppressing the activity of task irrelevant information (Yan et al., 2014). Other studies suggest that learning stems from better read-out mechanisms in higher-level visual areas (Law & Gold, 2009). Psychophysical studies have suggested that boosting sub-threshold, stimulus-related cortical activity can promote perceptual learning (Seitz & Dinse, 2007), with attention and reinforcement (provided by reward) increasing stimulus-related neuronal activity and facilitating learning (Ahissar, 2001; Pascucci et al., 2015; Seitz & Watanabe, 2005).

In parallel, increasing effort worldwide is being directed at applying visual perceptual training approaches to rehabilitate patients with various types of vision loss, including cortical blindness (CB), amblyopia (Huang et al., 2008; Levi & Li, 2009; Li et al., 2013; Li et al., 2011; Polat et al., 2004), macular degeneration (Baker et al., 2008; Kwon et al., 2012; Liu et al., 2007), myopia (Camilleri et al., 2014; Tan & Fong, 2008) and even keratoconus (Sabesan et al., 2017). Two critical factors that limit practical applications of VPL are: 1) the long duration of training usually required for adequate performance enhancement (e.g. in chronic CB patients, Huxlin et al., 2009), and 2) persistence of visual learning and/or recovered abilities after training ends. Non-invasive brain stimulation coupled with perceptual training has emerged as a potentially promising solution for both of these limitations in healthy adults (Ammann et al., 2016; Cappelletti et al., 2013; Chesters et al., 2017; Falcone et al., 2012; Fertonani et al., 2011; Mulquiney et al., 2011; Sehm et al., 2013; Snowball et al., 2013; Zoefel & Davis, 2017).

In CB, a form of vision loss caused by primary visual cortex (V1) damage, one approach shown to recover vision involves training on motion integration tasks in the blind field (Cavanaugh & Huxlin, 2017; Das et al., 2014; Huxlin et al., 2009; Melnick et al., 2016; Vaina et al., 2014). However, the training required to restore normal performance on this task in the blind field of CB patients typically involves months of daily practice, and is thus difficult to attain and sustain. As such, this represents an ideal task with which to ask whether non-invasive brain stimulation of early visual cortex during training can enhance and speed up the resultant perceptual learning. We tested this hypothesis first in visually-intact controls, and then, we conducted a follow up study in a small cohort of chronic CB patients.

We used two different forms of direct current stimulation to modulate cortical functioning and boost performance during learning (Fertonani & Miniussi, 2016): transcranial random noise (tRNS) and anodal direct current stimulation (a-tDCS). tRNS is an alternating current stimulation delivered at random frequencies within a defined frequency range, while a-tDCS uses constant, low positive (anodal) direct current to modulate cortical activity.

tRNS was first shown to enhance cortical excitability in the motor cortex (Terney et al., 2008) and subsequent studies reported that it can also improve perceptual functions when delivered over the visual cortex (Camilleri et al., 2016; Campana et al., 2014; Pirulli et al., 2013). Importantly, improvements attained with tRNS were seen within short periods of time and appeared to be long lasting, indicating a good potential for tRNS to promote clinically-meaningful, long-term plasticity (Tyler et al., 2018; van der Groen et al., 2018).

The second form of brain stimulation tested in the present experiments was tDCS. In motor cortex stimulation, tDCS is thought to increase cortical excitability near the anode and decrease excitability near the cathode (Miniussi et al., 2013). However, this excitation-inhibition model may be overly simplistic for other brain regions, including visual areas, and contradictory results have indeed been reported for the impact of tDCS on visual functions (reviewed in Miniussi & Ruzzoli, 2013). Still, there is evidence that tDCS might lead to vision improvements in amblyopia (Ding et al., 2016; Spiegel et al., 2013),

In summary, the present experiments asked if and what type of brain stimulation (tRNS, a-tDCS or both) could improve visual learning when administered during training in visually-intact humans and whether these improvements persist long after stimulation and training end. We then examined the translational potential of this approach to promote visual recovery in chronic CB patients. The motion integration task was chosen because it uses a stimulus whose detectability can be manipulated by changing its signal-to-noise ratio in the motion direction dimension (Figure 1). Early visual areas of the brain were targeted for stimulation because of their apparent role in mediating training-induced visual plasticity in physiological, imaging and brain stimulation studies (Barbot et al., 2018; Camilleri et al., 2016; Gratton et al., 2017; Kang et al., 2014; Rokem & Silver, 2010; Schwartz et al., 2002; Yang & Maunsell, 2003).

**Figure 1.**
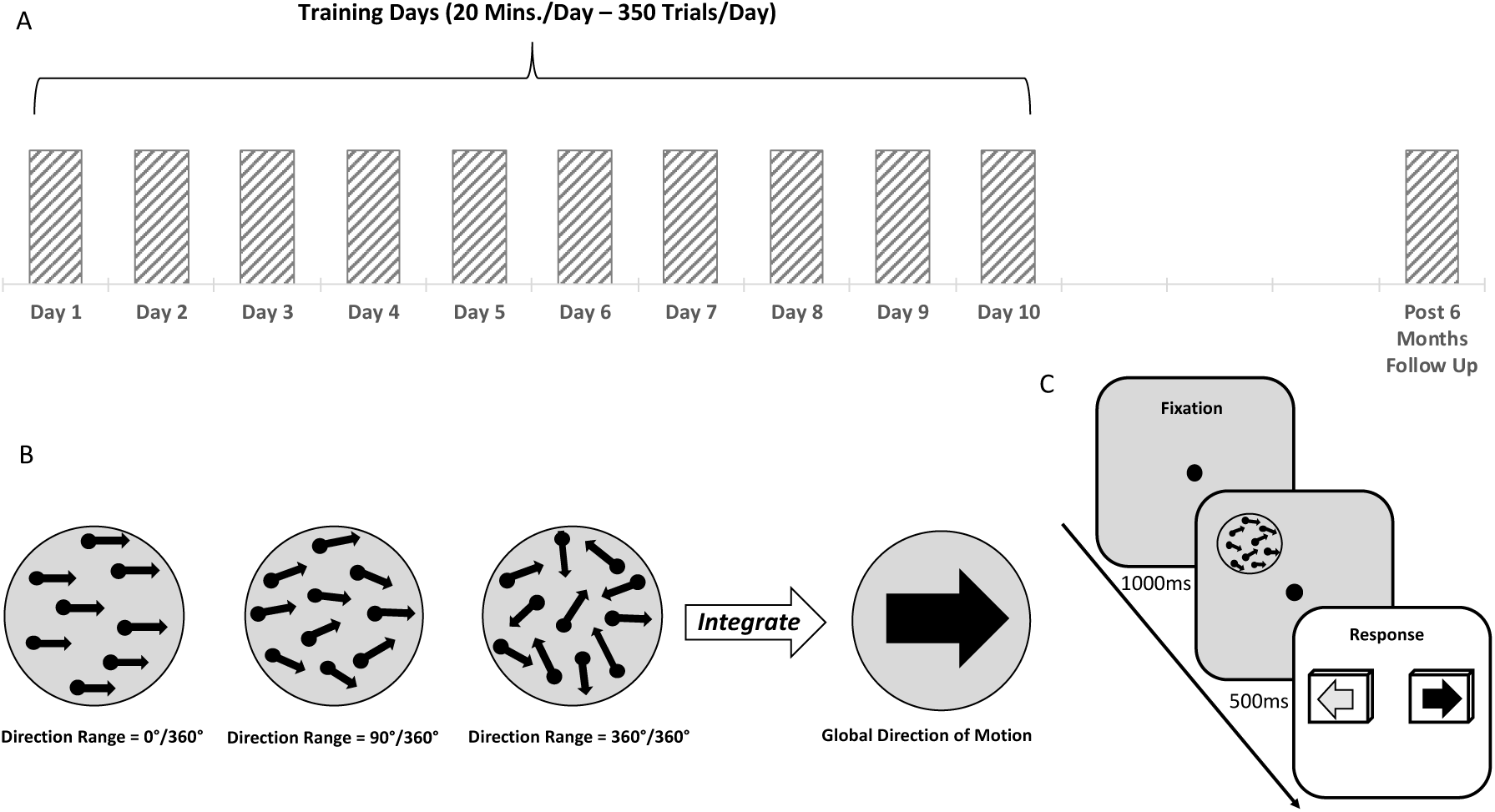
Experimental procedure and behavioral task. **A**. All participants were tested on a motion integration task to determine baseline performance in the first session (Day 1). They then underwent 9 days of training with or without online brain stimulation (Days 2 −10). Behavioral testing was performed again 6 months after the end of training/stimulation (Post 6 Months Follow Up). **B**. Example of stimuli with different direction ranges (0°, 90°, and 360°) used for the motion integration task, target dots were embedded in noise dots that are not shown in figure for clarity purposes (see Methods section for details). Normalized direction range (NDR, see text for details) 0 indicates fully random motion directions (360° range), while NDR = 100 indicates all signal dots moving in one direction (0° range). A two-alternative forced choice, adaptive staircase procedure was used to estimate the largest range of dot directions that subjects could correctly integrate to discriminate the global motion direction (leftward vs. rightward). **C**. Trial sequence used for training and to measure left-right motion discrimination thresholds. First, subjects were asked to fixate the central cross for 1000ms, immediately followed by a tone signaling the appearance of the stimulus, which was presented for 500ms. Subjects had to indicate the perceived global motion direction by pressing the left or right arrow key on the keyboard.

## Results

### Impact of pairing brain stimulation with training in visually-intact subjects

#### Learning of motion integration in control groups

Subjects recruited for the present experiments were divided into five training groups. Two experimental groups (bilateral tRNS and anodal stimulation) received stimulation over early visual areas. Their results were compared to three control groups: bilateral tRNS over parietal cortex, a sham control, and a no-stimulation control. Prior to the onset of training, there were no significant differences in normalized direction range (NDR) thresholds, a measure of direction integration performance, between the five training groups (*F_4,40_=1.15, p=0.35*). This confirmed that all five groups started with relatively similar levels of performance. As expected, all five groups benefited from perceptual training—for each group, performance at Day 10 was better than at Day 1 (all *t_8_ > 2.7*, all *p < 0.027*). This result is consistent with well-established effects of training on visual perception (Levi & Shaked, 2016; Watanabe, & Sasaki 2015). However, no significant differences in learning were observed between three of the groups — no-stimulation (training only), sham-stimulation + training and tRNS over parietal cortex + training. The lack of difference was observed irrespective of whether learning was expressed as a raw change in thresholds (NDR_Day1_ − NDR_Day10_; *F_2,24_ = 1.58, p = 0.23*), a percent change in thresholds ((NDR_Day1_ − NDR_Day10_)/NDR_Day1_; *F_2, 24_ = 2.93, p = 0.072*) or learning speed (linear regression slope; *F_2, 24_ = 2.36, p = 0.12*). To minimize the number of multiple comparisons between experimental and control groups, data from these three control groups were thus combined into a single, control data set for all subsequent analyses.

#### tRNS, but not a-tDCS, enhances learning

Comparison of the control data set with tRNS + training and a-tDCS + training revealed large differences in learning (Figure 2A-B). In addition to the expected main effect of training day (*F_3.4,144.1_ = 34.7, p = 10^−18^*), we found a main effect of group (*F_2,42_ = 3.35, p = 0.045*) and, notably, a significant group by day interaction (*F_6.9,144.1_ = 4.01, p = 0.01*). As suggested by this significant interaction, the amount of learning differed among the three groups (Figure 2A; *F_2,42_ 9.39, p 0.0004*). Specifically, tRNS + training induced stronger learning than both the combined control group (*p* = *0.002*) and a-tDCS + training (*p* = *0.001;* all post-hoc tests are Tukey HSD), whereas a-tDCS outcomes did not differ significantly from those attained by the combined control group (*p = 0.53*). The same pattern of results was observed when we considered group differences in terms of percent improvement from pre- to post-test (Figure 2B, *F_2,42_=10.8, p=0.00016*). Again, tRNS + training resulted in larger percent improvement than attained by controls (*p=0.001*) and a-tDCS + training (*p=0.0002*). Once again, performance following a-tDCS + training did not differ significantly from that in the combined control group (*p* = *0.28*). In all groups, learning was well described by a linear trend (Figure 2A). Slopes, however, differed among groups (Figure 2C; *F2,42=7.8, p=0.001*), with tRNS + training generating faster learning than in the combined control group (*p=0.008*) or a-tDCS + training group (*p=0.001*). In contrast, a-tDCS did not induce significantly faster learning than attained by the combined control group (*p=0.30*).

**Figure 2.**
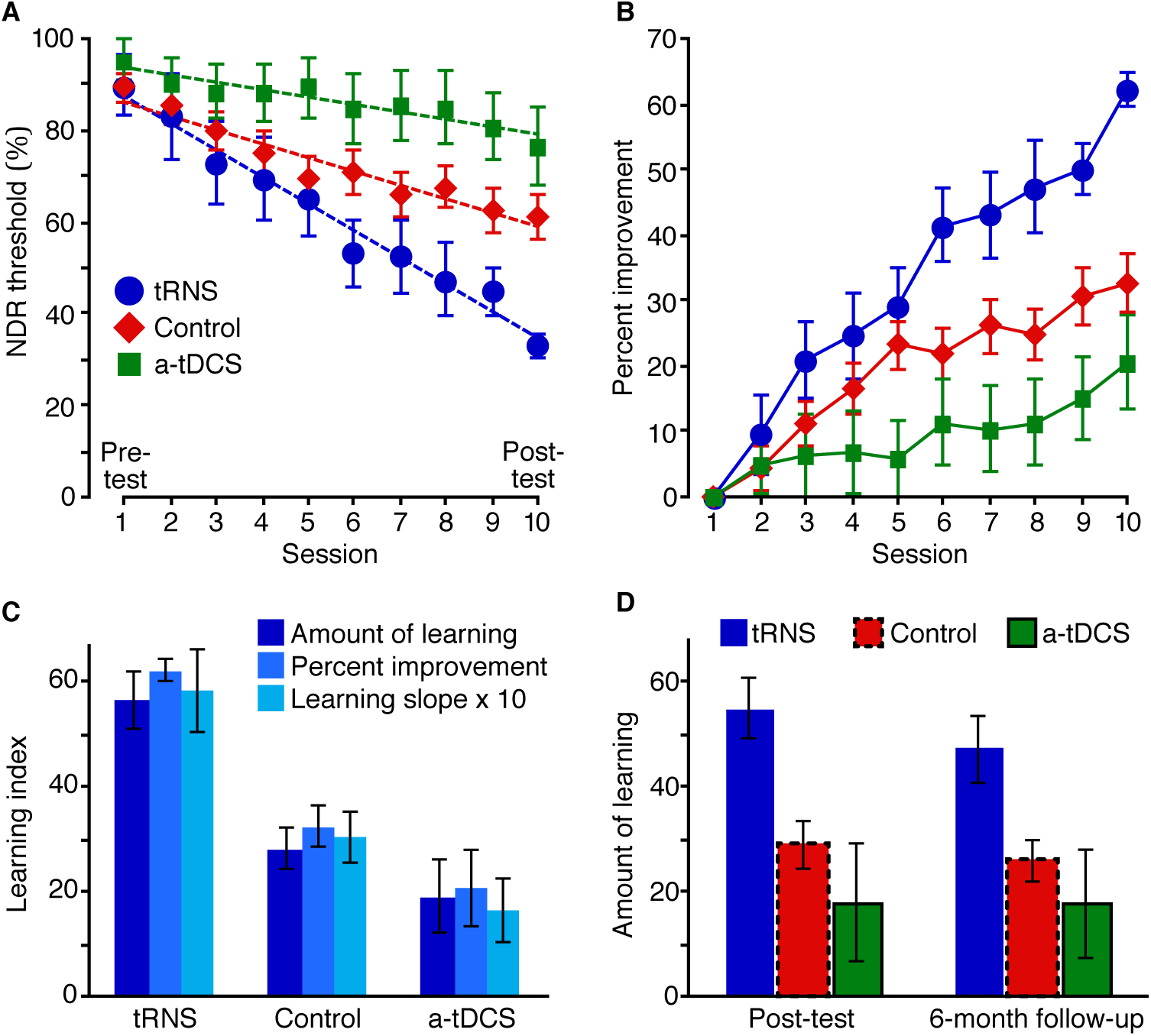
Effects of brain stimulation on perceptual learning in visually-intact subjects. **A**. Normalized direction range (NDR) thresholds for the control groups, tRNS and a-tDCS. Dashed lines are linear fits, indicating the learning slope. **B**. Same data as in A, but expressed as percent improvement relative to Day 1 thresholds. **C**. Learning index computed in three different ways. tRNS group exhibited a significantly stronger amount of learning (Day 1 – Day 10; F_2, 42_ = 9.39, p = 0.0004; all Tukey HSD p < 0.002), percent improvement (100*(Day 1 – Day 10)/Day 1; F_2, 42_ = 10.8, p = 0.00016; all Tukey HSD p < 0.001) and learning slope (F_2, 42_ = 7.8, p = 0.001; all Tukey HSD p < 0.008) than both the control and a-tDCS groups. **D**. Amount of learning, defined as the difference from Day 1 thresholds, at the end of the training (left) and 6 months after (right). Error bars are ±1 SEM.

As tRNS administered during training appeared to cause faster learning, we analyzed at what time point the tRNS group began to diverge from the other two groups. This occurred on Day 6 (*F_2,42_=5.03*, *p=0.01*), at which point, tRNS showed stronger learning than both the combined control group (*p=0.03;* Tukey HSD) and a-tDCS + training (*p=0.01;* Tukey HSD).

In sum, we found strong evidence for enhanced learning in the tRNS + training group, with faster learning than both the combined control group and a-tDCS + training. As detailed above, this finding was supported irrespective of how learning was defined. While it may seem that a-tDCS, as administered in our study, might actually hinder learning (Figure 2A-B), this effect was not statistically significant.

#### Persistence of stimulation-enhanced perceptual learning

Next, we asked whether the observed enhancement of perceptual learning by tRNS remained stable over an extended period of time. To address this question, we re-tested participants 6 months after completing the 10-days of training with and without the different forms of stimulation. The post-test, however, was performed *<u>without</u>* brain stimulation. The subjects re-tested at 6 months included 37 of 45 original participants (*n=8* for tRNS + training group, *n=22* for the combined control group, and *n=7* for a-tDCS + training group). Figure 2D contrasts the amount of learning at Day 10 (NDR_Day 1 –_ NDR_Day 10_) with that exhibited 6 months after the end of training (NDR_Day 1 –_ NDR_6-months_). There was a small, non-significant loss in performance for the three groups (no main effect of testing day; *F_1,34_ = 3.32, p = 0.8*) and no interaction (*F_2,34_ = .88, p = 0.43*). We only found a main effect of group, confirming that the group differences at the end of 10 days of training remained unaltered 6 months after training (*F_2,34_ =* 4.68, *p = 0.0*2). Thus, it appears that tRNS enhanced perceptual learning over the long-term - at least 6 months after the end of both training and brain stimulation. Moreover, this persistent enhancement was observed without brain stimulation at the 6 months follow-up. This suggested that the enhancement of perceptual learning by tRNS was not due to online or short-term optimization of visual processing, but instead, resulted in consolidated sensory learning.

##### tRNS boosts training-induced visual recovery in cortically-blind patients

Given our finding that tRNS, but not a-tDCS, over occipital cortex considerably enhances perceptual learning in neurotypical subjects, we next asked whether tRNS is also able to enhance training-induced visual recovery in chronic, V1-damaged patients with CB. To the best of our knowledge, tRNS has not been attempted in brain-damaged patients. Moreover, whether tRNS over early visual areas could enhance learning in CB patients is an open question, as learning in this patient population can exhibit properties not found in neurotypical subjects (e.g. Cavanaugh & Huxlin, 2017; Das et al., 2014; Vaina et al., 2014), and since by definition, part of early visual cortex that would normally be stimulated is damaged. Hence, we sought to perform a preliminary, proof-of-concept study in five patients with occipital damage resulting in homonymous visual field defects measured with visual perimetry (Table 1, Figure 3).

**Table 1.** Demographic data for CB patients. Visual fields defects were assessed with automated perimetry. The last column indicates the time between stroke and in-lab testing. Patient RNS3 had a traumatic brain injury, while all other patients suffered strokes.

| Patient | Sex | Age at Testing | Visual Defect | Lesion | Time Since Lesion (Months) |
| --- | --- | --- | --- | --- | --- |
| RNS1 | F | 59 | Left lateral homonymous hemianopia | Stroke affecting right fronto-parieto-occipital lobe | 36 |
| RNS2 | M | 66 | Left lateral homonymous quadrantanopia | Stroke affecting right posterior capsule-thalamic and occipital lobe | 15 |
| RNS3 | M | 53 | Bilateral upper homonymous quadrantanopia | Traumatic brain injury affecting Left fronto-temporo-parietal region | 108 |
| Sham1 | M | 69 | Right lateral homonymous hemianopia | Stroke affecting left occipital lobe and left posterior capsule-thalamic area | 24 |
| Sham2 | M | 72 | Right lateral homonymous quadrantanopia | Stroke affecting left occipital lobe and internal capsule | 9 |
| U1 | F | 63 | Right homonymous hemianopia | Stroke affecting left occipital lobe | 10 |
| U2 | F | 67 | Left lower homonymous quadrantanopia | Stroke (hemorrhagic) affecting right occipital lobe | 26 |
| U3 | F | 54 | Left homonymous hemianopia | Stroke affecting right occipital cortex | 7 |
| U4 | M | 47 | Bilateral hemianopia | Bilateral stroke damage affecting occipital lobes | 16 |
| U5 | M | 53 | Left homonymous hemianopia | Stroke affecting right occipital lobe | 12 |
| U6 | F | 44 | Left homonymous hemianopia | Stroke affecting right occipital lobe | 2.5 |

Visual perimetry was used to identify the blind field borders and select training locations in the blind field (Figure 3). We randomly assigned five patients from our Italian study site to either tRNS + training (*n=3, RNS1-3*) or sham stimulation + training (*n=2, Sham1-2*). Data from an additional six CB patients who trained identically, but without brain stimulation (unstimulated, U1-6), at our United States study site, were also analyzed for comparison. All patients, at both study sites, trained for 10 days using random dot stimuli, as in neurotypical subjects (Figure 1). Because patients have difficulty seeing motion, their stimuli, unlike those for neurotypical subjects, did not include noise dots.

**Figure 3.**
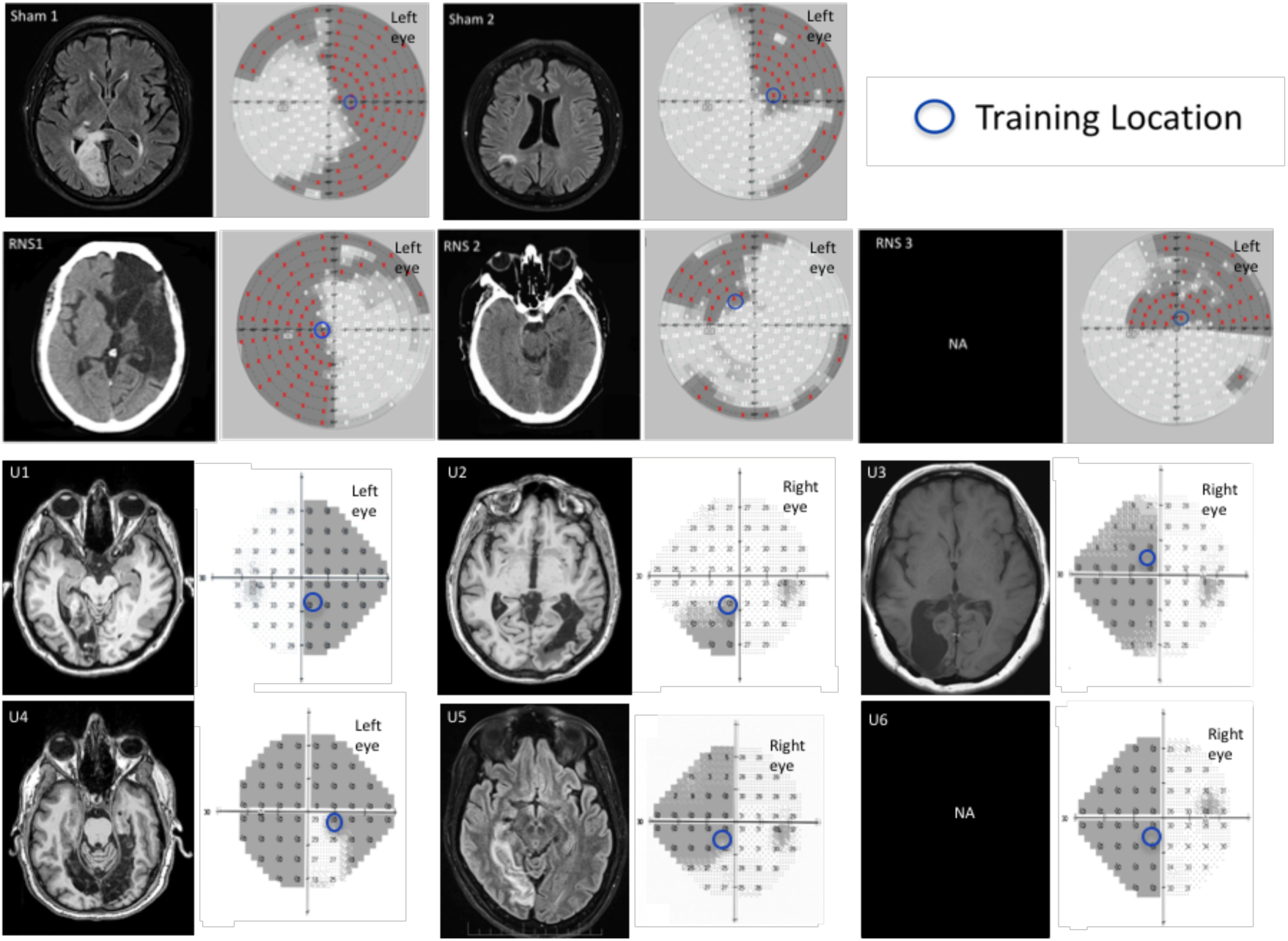
Neuroradiological images and Visual Field Perimetries of chronic CB patients. All patients sustained damage of early visual areas or the optic radiations resulting in homonymous visual field defects as shown by the visual field perimetries, next to each brain image. Within the perimetry images (patients in top two rows: Sham1, Sham2, RNS1, RNS2 and RNS3): red marks and shading areas indicate the patients’ blind field. Bottom two rows: composite Humphrey visual field maps are reported for each unstimulated patient, superimposed shading indicates the blind field. Numbers indicate the luminance detection sensitivity in the given position expressed in decibels for all perimetries. For all patients, the blue circles indicate the location where we positioned the training stimuli and the size of the blue circles indicates the size of the training stimulus (see *“Global direction discrimination testing and training in patients”* in the Methods section for details). Radiological images were not available for patients RNS3 and U6.

As expected (Huxlin et al., 2009; Das et al., 2014; Cavanaugh et al., 2015; Cavanaugh and Huxlin 2017), all patients performed considerably worse on global motion integration in their blind field compared to neurotypical subjects. This was the case even at the easiest stimulus level (NDR = 100, with all dots moving in the same direction), where none of the patients exhibited ceiling level performance in their blind field. As such, we used percent correct at 100 NDR as the measure of performance (see Materials and Methods for details). Sham-stimulated patients exhibited no significant change in performance across the 10 days of training (Figure 4A) as evidenced by learning slopes that were not significantly different from 0 (all *p>0.63;* see Methods for bootstrap procedure used to analyze data from individual patients). This was comparable to the lack of learning observed in the six unstimulated patients, who also did not exhibit significant learning slope over their first 10 days of training (Figure 4C; all *p* > 0.13). In contrast, tRNS coupled with training enhanced the rate of global motion discrimination learning in CB patients (Figure 4B), who exhibited significantly positive learning slopes (all *p<0.0048*). We also examined the change in performance from Days 1–2 to Days 9–10, averaging results over two consecutive days to minimize the effects of day-by-day fluctuations. The analysis showed significant change only for patients trained with tRNS (Figure 4D; tRNS: all *p < 0.0002;* Sham: all *p > 0.24;* Unstimulated: all *p > 0.080*).

**Figure 4.**
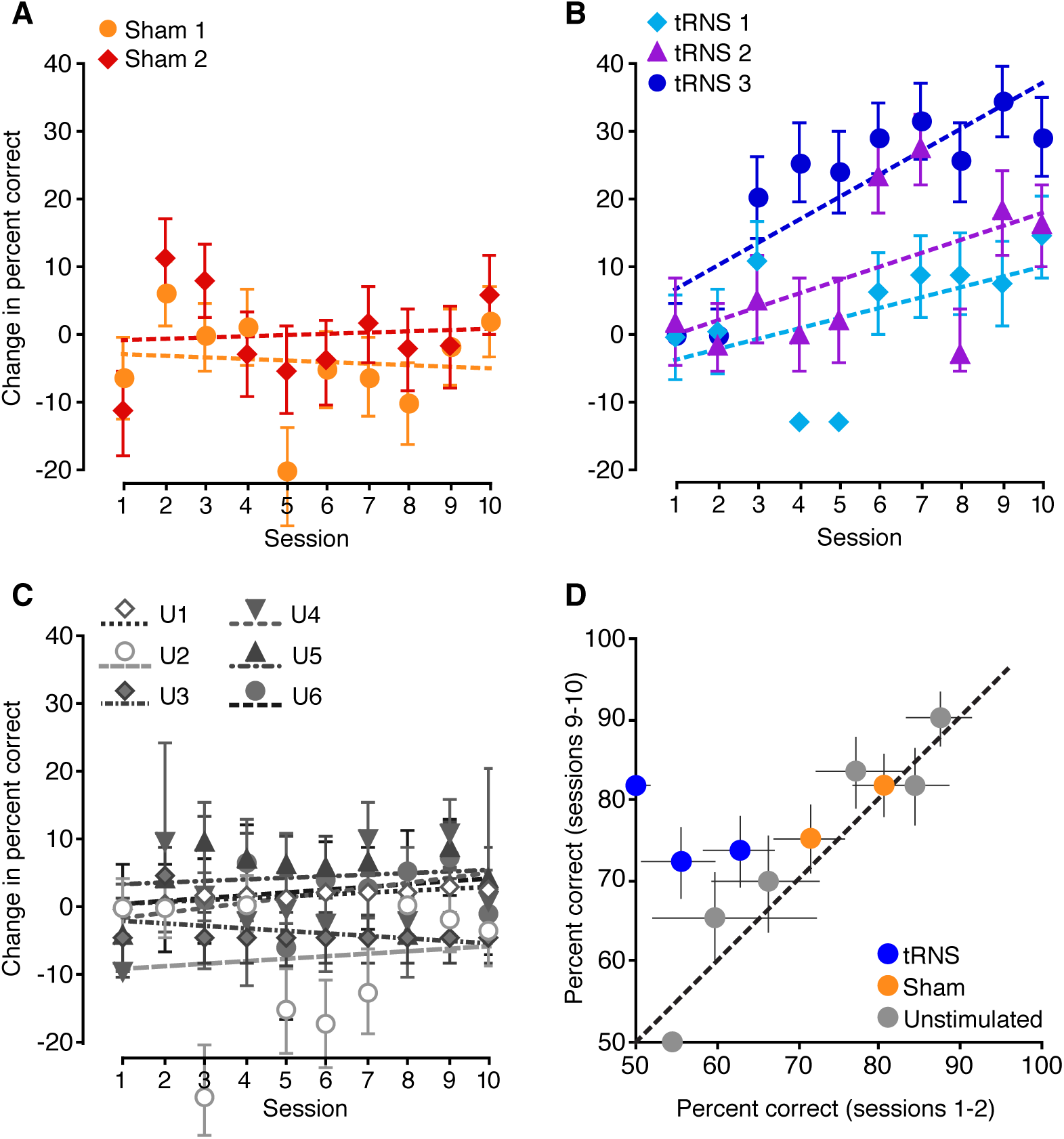
The effects of brain stimulation on perceptual learning in CB patients. Task performance over 10 training days for patients who underwent (**A**) sham stimulation, those (**B**) who received tRNS, and (**C**) six unstimulated patients. Raw percent correct performance was normalized by subtracting the average percent correct for the first two training days. **D**. Comparison of raw percent correct averaged over the first two days against raw percent correct for the final two training days. Significant learning was observed only for patients who trained with tRNS. All error bars= 95% confidence intervals. For panels A-C, all lines are linear fits, indicating learning slope.

## Discussion

In the present study, we asked whether brain stimulation over early visual cortex could be used to boost and speed up visual perceptual learning in healthy humans, and those affected by cortically-induced blindness. In healthy humans, we also asked if enhancements attained from training coupled with stimulation persisted over long periods of time. Our results show that tRNS applied bilaterally over healthy, early visual cortex speeds-up and boosts performance gains during visual perceptual learning. Over 10 days, neurotypical subjects in the tRNS + training group exhibited about a 60% improvement in motion integration thresholds (Figure 2B), which was two and three times as strong as learning attained by the control and a-tDCS groups, respectively. This finding was supported irrespective of whether learning was defined as a raw change in NDR threshold, a percent improvement or the slope of a linear fit to the data. The observed benefit of tRNS over training alone or sham stimulation + training, or tDCS over parietal cortex is consistent with evidence that tRNS is especially effective at promoting plasticity when coupled with a relevant stimulus, and when it is applied over relevant [in our case, occipital] brain areas (Cappelletti et al., 2013). In contrast, we found no such benefit of a-tDCS over occipital cortex. Finally, we provide for the first time preliminary evidence that tRNS enhances vision recovery in CB patients with V1 damage. Moreover, we demonstrate seemingly safe usage of this technique in a class of stroke patients, with no side-effects reported. Notably, with respect to training-induced recovery, tRNS enabled improvements in visual task performance of chronic CB patients in their blind field, over a tiny fraction of training days typically required to induce such improvements in the absence of brain stimulation (on average, 72 to 80 training days are required to recover global direction discrimination performance at a given blind field location – see Cavanaugh & Huxlin, 2017; Das et al., 2014; Das & Huxlin, 2010; Huxlin et al., 2009; Melnick et al., 2016). Indeed, we observed a steady and significant increase in performance for three patients trained with tRNS over 10 days, with no such effects in patients that trained with sham stimulation. Chronic CB patients are a population that would especially benefit from enhanced perceptual learning because vision recovery using conventional training methods usually takes many months of daily training (Cavanaugh & Huxlin, 2017; Das et al., 2014; Das & Huxlin, 2010; Huxlin et al., 2009; Melnick et al., 2015).

Interestingly, while the position of the stimulating electrodes corresponded to early visual areas that included V1, improvement occurred despite V1 being damaged in the CB patients. This might suggest that tRNS’ neuromodulatory benefit likely impact any residual V1, but also adjacent visual areas such as V2 and V3, which might have also supported visual learning as indicated in imaging studies in CB patients (Barbot et al., 2018; Ajina et al., 2015; Martin et al., 2012; Raemaekers et al., 2011). V2 and V3 are usually spared in CB patients and they might have played a pivotal role in supporting recovery during training. Interestingly, physiological studies have shown that V2/V3 inactivation may degrade cortical motion sensitive areas’ ability to work efficiently (Ponce et al., 2011), while they might support learning when they function normally (Law & Gold, 2008). In summary, while plasticity of spared visual circuits is generally believed to underlie visual recovery, the specific neural mechanisms involved remain unclear.

An additional and important finding in the present study was the observation that enhancement of perceptual learning induced by tRNS can persist over an extended period of time (6 months in the present study) beyond the end of stimulation and training. This is important since stimulation-enhanced perceptual learning would have limited practical use if its beneficial effects diminished over time. Additionally, because all subjects performed the behavioral task with no stimulation at the follow-up, 6 months timepoint, we infer that the benefits of tRNS go beyond an online enhancement of visual processing and likely involve plastic changes that persist within the visual system, allowing it to more effectively process global motion stimuli. As such, we conclude that consolidation of learning occurred in our subjects. The persistence effects observed here are particularly notable, as many VPL studies failed to see long lasting effects and/or transfer of learning to other tasks (Dosher & Lu, 2017).

Some important questions arising from the present results are: 1) how does brain stimulation enhance the effects of perceptual learning? and 2) why do some forms of stimulation prove effective and others not? Anodal tDCS did not exert a beneficial effect in our study - a surprising result given results from previous studies stimulating the early visual cortex (Antal et al., 2004). One possible explanation is that the strongest effects of tDCS were reported offline, where a-tDCS was delivered *prior* to the measured behavior (Pirulli et al., 2013). Another possibility is that a-tDCS is not the ideal neuromodulatory technique for repeated sessions. While it alters membrane potentials and hence exerts increased excitability, it may also engage inhibitory homeostatic mechanisms during repeated sessions (Fertonani et al., 2011; Peters et al., 2013).

Our observation that visual enhancements persist long after both training and tRNS ended does constrain possible mechanistic explanations. Multiple types of transcranial electrical stimulation have been shown to alter excitability in cortex, and the longer time course of direct current stimulation effects has been suggested to relate to homeostatic changes in membrane potential (Ardolino et al., 2005; Liebetanz et al., 2002; Nitsche et al., 2003; Terney et al., 2008) or gate threshold (Bikson & Rahman, 2013). However, direct evidence that transcranial stimulation alters the dynamics of networks known to be related to perceptual learning, such as dopaminergic reward networks (Pascucci et al., 2015; Seitz & Watanabe, 2005), has not yet been provided. All we can state here is that tRNS does not appear to globally affect reward networks, as there was no boost in visual performance or learning seen from stimulation over parietal cortex.

An interesting hypothesis is that tRNS-related visual performance improvements might derive from the state of the neurons at the time of stimulation (Silvanto et al., 2008), and that adding noise to the cortex might enhance sensory detection, in particular when stimuli are presented at threshold and embedded in noise (Abrahamyan et al., 2015). Several short-term mechanisms have also been proposed to explain the effects of tRNS, and a favored hypothesis involves stochastic resonance, whereby random frequency stimulation in tRNS appears to boost responses of neural populations to weak inputs (Moss et al., 2004; Pirulli et al., 2013; Miniussi & Ruzzoli, 2013; Schwarzkopf et al., 2011; van der Groen & Wenderoth, 2016; van der Groen et al., 2018; Herpich et al., 2018). An alternative hypothesis proposes temporal summation of excitatory signals between visual stimulation and electrical stimulation (Fertonani et al., 2011; Pirulli et al., 2013; Terney et al., 2008), and selective enhancement of active neural networks (Bikson et al., 2013; Fertonani & Miniussi, 2016; Luft, 2014; Miniussi et al., 2013). It is important to note that one or more of these short-term mechanisms may be the first step in a longer-term cascade that results in persistence of learning. For instance, stronger activation of task-relevant neurons due to temporal summation or stochastic resonance may encourage a shift towards greater plasticity in sensory processing and/or readout. However, the time course of effects observed in the present study, and especially their persistence, suggests that online phenomena (i.e. during stimulation or shortly thereafter) are not the only ones at play with respect to learning enhancements induced by tRNS. Interestingly, studies on perceptual learning in animal models have shown that learning might boost the modulation in neuronal tuning to stimulus components relevant to the task (Liu & Pack, 2017). If learning was associated with changes in the tuning characteristics of neurons (for a review see (Gilbert & Li, 2012)), we could speculate that tRNS coupled with behavioral training might facilitate and consolidate this plastic change, which could then persist across months (Snowball et al., 2013).

Regardless of its precise mechanism of action, here we provide empirical evidence for the potential usefulness of tRNS coupled with visual training on a patient population that requires perceptual learning in order to attain visual recovery. V1-damaged patients with chronic CB are able to recover some visual abilities within their scotoma, but only after intensive and repetitive training over many months of daily practice (Das et al., 2014; Huxlin et al., 2009). The application of safe, painless neurostimulation in situations like this, where perceptual learning is directly proportional to the quantity of vision recovered (Cavanaugh & Huxlin, 2017) has the potential to dramatically improve quality of life and treatment outcomes (Cavanaugh et al., 2016). Therefore, results from our experiment with CB patients suggest that tRNS might be a viable adjunct procedure to speed up the recovery process. Remarkably, even though the physiological effects of tRNS upon the damaged early visual cortex are currently unknown, our data show that tRNS can help overcome reduced and/or partially absent functionality and boost learning in the blind field.

## Conclusion

Perceptual training administered concurrently with tRNS for 10 days in healthy subjects achieves large and lasting vision improvement in the perception and processing of global motion direction. That tRNS together with visual training also enhances learning in chronic, V1-damaged patients with CB, suggests that stimulation can be effective even if it is delivered over partially damaged cortex, long after the damage has occurred. Based on these findings, the efficacy of visual training and rehabilitation procedures could be dramatically increased by incorporating specific forms of non-invasive, transcranial electrical neuromodulation, both in healthy and neurological populations.

## Materials and Methods

### Regulatory approval

The present study was approved by the ethical committee of the University of Trento, the ethical committee for clinical experimentation of the “Azienda Provinciale per i Servizi Sanitari” (APSS) and by the Institutional Review Board of the University of Rochester Medical Center, Rochester, NY, USA. The work was conducted after obtaining written, informed consent from each participant.

### Visually-intact participants

A total of 45 subjects participated in the experiment (mean age: 19.9 years old; range: 19–36; 32 females and 13 males). All subjects were right-handed, neurologically normal, with normal or corrected-to-normal vision and gave written, informed consent prior to the beginning of the study, according to the ethical standards of the Declaration of Helsinki.

### Cortically-blind patients

Eleven patients participated in the study: five patients recruited at the Center for Neurocognitive Rehabilitation (CeRiN) affiliated to the University of Trento and the Rehabilitation Hospital “Villa Rosa”, in Pergine, Italy and six patients recruited at the Flaum Eye Institute of the University of Rochester Medical Center in Rochester, NY, USA. Patients were recruited 2.5–108 months after damage to their early visual areas (Table 1, median: 15 months), causing homonymous visual defects, confirmed by neurological reports, neuro-radiological exams and automated visual perimetry (Optopol PTS 1000 Visual Field, Canon or Humphrey Field Analyzer HFA II 750, Carl Zeiss Meditec). The patients gave written, informed consent at their respective study site before participating.

Ten of the patients (RNS1-2, Sham1-2, U1-6) suffered from stroke involving the territory of the posterior cerebral artery, as confirmed by radiological examinations and reports (Figure 3). One patient (RNS3) suffered from traumatic brain injury. Although from the neuro-radiological report, V1 was not directly affected by trauma, there were indications of visual fields defects and his visual perimetry showed a clear, homonymous, bilateral upper quadrantanopia; hence, we decided to enroll him in the training procedure (note that data for each patient were computed and shown individually). None of the patients had history or evidence of degenerative or psychiatric disorders. All participants were right handed, with normal or corrected-to-normal visual acuity and none exhibited visual or other forms of neglect, as determined by neurological examination.

### Study design

Visually-intact controls were randomly assigned to one of five groups. This included two experimental groups. In the first group, tRNS was delivered over early visual areas (electrodes positioned bilaterally, centered over O1 and O2 of the EEG system coordinates, for the left and right hemisphere respectively). In the second group, anodal transcranial direct current stimulation (a-tDCS) was delivered over the occipital cortex (the anode and the cathode were positioned over Oz and Cz, respectively). While we used bilateral occipital montage for the tRNS condition to match the positioning of other successful studies that found improved performance with tRNS and likely increased excitability in the visual cortex (Romanska et al., 2015; Herpich et al., 2018) particularly with motion discrimination tasks (van der Groen et al., 2018), we chose unilateral montage for the a-tDCS condition, the optimal montage to increase cortical excitability with anodal stimulation of the visual cortex (Antal et al., 2004). Stimulation was concurrent with the training task. There were also three control groups: a sham control, a no-stimulation control, and an active control where bilateral tRNS was applied over parietal cortex (over P3 and P4, regions likely involved in but not critical for global motion discrimination (Battelli et al., 2001; Greenlee & Smith, 1997)). Over 10 days, all neurotypical subjects were trained to discriminate the left or right, global direction of random dot motion stimuli (350 trials/session/day). Day 1 was considered the pre-training session, while Day 10 was used as the post-training session. Finally, a long-term follow-up was performed six months after the post-training session. During this follow-up, participants repeated the behavioral baseline tests. Critically, no stimulation was delivered at this time.

All cortically blind patients underwent 10 days of training, following the same Day 1 to Day 10 procedures as neurotypical subjects (Figure 1). Patients in Italy were randomly assigned to one of two experimental groups. Three patients received tRNS over early visual areas during training, whereas two patients received sham stimulation during training, with electrodes placed on the same location as for tRNS. Patients in the USA underwent global direction discrimination training without brain stimulation.

### Apparatus and procedures

Unless otherwise stated, we used methods and apparatus in neurotypical subjects and patients that were as similar as possible (see below), in Italy and the USA. For subjects who underwent brain stimulation, all experimental days took place in the same room, under the same light and noise conditions, and with the same apparatus. During each session, participants were positioned on a chinrest-forehead bar combination to stabilize their heads, and to place their eyes 57 cm from the stimulus-presenting computer monitor. Visual stimuli were generated on a MacBook Pro running software based on the Psychophysics Toolbox (Brainard, 1997; Pelli, 1997) in Matlab (*MathWorks*). Stimuli were presented on a linearized SensEye 3 LED 24 Inch (*BenQ*) monitor with a refresh rate of 120 Hz for the patients tested in Italy and a CRT monitor (HP 7217 A, 48.5 × 31.5 cm, 1024 × 640 pixel resolution, 120 Hz frame rate) for the patients in the USA. The LED monitor was luminance-calibrated with gamma=1 using a professional monitor calibrator (Datacolor Spyder 5), while the CRT monitor was calibrated with a ColorCal II automatic calibration system (Cambridge Research Systems). Eye fixation for all subjects was controlled in real time using an EyeLink 1000 Plus Eye Tracking System (*SR Research Ltd., Canada*) whose infrared camera monitored the pupil center and corneal reflection of the left eye. Limits were set so that if the participant’s eye moved > 1.5° in any direction away from the fixation spot during stimulus presentation, loud tones sounded and the currently displayed trial was aborted and excluded from the final analysis.

#### Global direction discrimination testing and training in neurotypical subjects

We first measured direction range thresholds for left-right motion discrimination of circular stimuli that contained a limited percentage of signal dots (Newsome & Pare, 1988, Huxlin & Pasternak, 2004; Levi et al., 2015) and were centered at [-5, 5] deg in the visual periphery. To match initial task difficulty across observers, motion coherence (Newsome & Pare, 1988) was calibrated for each subject individually, as previously reported (Levi et al., 2015). The motion coherence of the stimulus was chosen based on preliminary testing aimed to identify a motion signal level that allowed participants to perform the discrimination task just above chance (50% correct). For all but 3 subjects, random dot stimuli contained 40% motion signal. Three subjects were trained with a stimulus containing 30% coherent motion, to allow enough room for improvement. Once a motion signal level was selected for each participant, the task used a QUEST adaptive staircase (Watson & Pelli, 1983) to estimate the broadest distribution of dot directions that subjects could correctly integrate to discriminate the global direction of motion as leftward or rightward. During training, task difficulty was adaptively modulated by adjusting direction range of signal dots (Huxlin & Pasternak, 2004) using twelve randomly interleaved 25-trial Quest staircases in each daily session. The resultant individual trials within each daily session were fit with Weibull functions (maximum likelihood fits). Thresholds reported in the paper, corresponding to 82% correct, were taken from these estimated Weibull functions and are reported as normalized direction range (NDR) thresholds, such that an NDR of 0% equals fully random motion (360° range of dot directions) and NDR of 100% indicates all signal dots moving in one direction (0° range). The random dot stimuli were presented within a circular aperture 5° in diameter at a density of 2.6 dots/deg^2^. Each dot had a diameter of 0.06° and moved at a speed of 10°/s with a lifetime of 250ms. Stimulus duration was 500ms. Each participant started training with direction range in the random dot stimulus set to 0°.

Trial sequence was as follows: participants were asked to fixate on a central cross for 1000ms, immediately followed by a tone signaling the appearance of the stimulus, which was presented for 500ms. Once the stimulus disappeared, participants had to indicate the perceived global direction of motion by pressing the left or right arrow keys on the keyboard. The two motion directions (leftward and rightward) were randomized across trials. Auditory feedback was provided indicating the correctness of the response on each trial.

During training, stimuli were presented monocularly to the left eye for 10 days (one session/day from Monday to Friday, for 2 consecutive weeks) while subjects received either active, sham or no stimulation. We chose monocular presentation to closely match the procedure used by Huxlin and colleagues (Huxlin et al., 2009) on patients and by Levi et al. (2015) on healthy participants using the same visual stimuli. Subjects performed 350 trials/day for a total of 3500 trials by the end of the 2 weeks of training. The total duration of the daily training session for each group was set to last about 20 min.

#### Global direction discrimination testing and training in patients

First, we spatially mapped motion discrimination performance in each patient to identify a blind field location where training should be performed. Here, we used the task described above for neurotypical subjects, with two adjustments to make the task easier for patients: coherence and NDR were set to 100% (the easiest possible settings) and the number of trials per training day was lowered to 250. Fixation was enforced, as in visually-intact subjects, and each trial was initiated by fixation of a small circle in the center of the screen. During mapping, stimuli were first presented in the intact field, at locations close to the border with the blind visual field, and patients performed 100 trials of the global direction discrimination task per location. This allowed us to ensure that each patient understood task demands and to assess normal baseline performance on an individual basis. Stimulus location was then moved progressively into the blind field, with 100 trials of the global direction discrimination task performed at each location, until global motion direction discrimination dropped below normal levels (measured in the intact field); this was selected as the training [blind field] location, and the whole stimulus was hence situated fully inside the perimetrically-defined blind field (blue circles in Figure 3).

For comparison purposes, we also included data from six patients trained in the Huxlin lab at the Flaum Eye Institute of the University of Rochester (USA) with the same behavioral protocol, but without any brain stimulation. Five out of six unstimulated patients, trained with 300 trials per training day, while one trained with 225 trials/day. Thus, on average, unstimulated patients completed 15% more trials than tRNS/sham-stimulated patients. This makes unstimulated patients a conservative comparison group.

On Day 1, patients performed considerably worse on global motion integration in their blind field compared to neurotypical subjects, even at the easiest stimulus level (NDR = 100%). On Day 10, no patient performed better than 85% correct at NDR of 100% (i.e., with all dots going in the same direction). Given this range of performance by patients, we chose to report percent correct at 100% NDR as the measure of performance. This allowed us to avoid issues with noisy thresholds estimates for sub-threshold performance, while still retaining a sufficient dynamic range to capture training-induced improvements in performance.

### Stimulation Protocols

tDCS and tRNS were delivered using a battery-driven stimulator (*DC-Stimulator-Plus, NeuroConn GmbH, Ilmenau, Germany*) through a pair of saline-soaked conductive rubber electrodes (35cm^2^). Each subject was randomly assigned to one of the 5 stimulation groups as described earlier. The electrodes were bilaterally placed over the target areas identified following the 10–20 EEG reference system. Subjects wore a Lycra swimmers’ cap to keep electrode in place, and we ensured that the skin and hair between the electrodes were completely dry, otherwise preventing the current from reaching the brain. The intensity of stimulation was set to 1.0mA, and was delivered for 20 min with a fade in/out period of 20 seconds. For the a-tDCS group the polarity of the active electrode was anodal. For the tRNS condition, the random noise stimulation was applied with a 0mA offset at frequencies of alternating current ranging from 101 to 640 Hz (high frequency tRNS). For the Sham stimulation group the stimulation (using the same electrode montage as in the tRNS condition) was shut down after 20 s. At the end of each session, we asked all subjects to fill out a questionnaire about potential discomfort or any unusual sensation they experienced during the stimulation. Only minor side-effects were reported by the tDCS group (2 subjects reported slight itching under the electrode, 1 subject reported a slight subjective temperature increase under the electrode), whereas none of the tRNS group participants reported any sensation of being stimulated.

## Data Analysis

The Shapiro-Wilk test was used to control for the normality of data distribution. Data Sphericity was addressed using Maulchy’s test, and Greenhouse-Geisser correction was used in case of non-sphericity of the data. Levene’s test was used to address the assumption of equality of variances. P-values were considered significant for values of < .05. To correct for multiple comparisons in post-hoc testing, we used Tukey HSD correction. The effect sizes are reported as the partial Eta-squared (ηp^2^) values.

To analyze data from individual patients, we performed the following bootstrap analysis. First, for each subject and each training day, we generated 10,000 bootstrap samples by selecting, with replacement, from the set of available individual trials. Then we fit Weibull functions to all 10,000 samples, and from resultant fits, we computed percent correct performance at 100% NDR (the easiest difficulty level). Because most of the individual trials for patients were collected near 100% NDR, these percent correct estimates were more robust than threshold estimates, which in many cases were estimated to be higher than 100% NDR. Thus, for each training day, we had 10,000 estimates for each patient’s percent correct performance, allowing us to estimate 95% confidence intervals (Figure 4). From this set of estimates, we created 100,000 full data sets for each patient (random sampling with replacement). This allowed us to estimate p-values for learning slope and amount of learning analyses reported in the Results. For the slope analysis, we simply computed the proportion of data sets that had negative learning slopes, multiplying results by 2 to get two-tailed p-values. For the amount of learning analysis, we computed the proportion of data sets where Day 1–2 performance was better than Day 9–10 performance, multiplying results by 2 to get two-tailed p-values.

## Acknowledgements

The present study was funded by the Autonomous Province of Trento, Call “Grandi Progetti 2012”, project “Characterizing and improving brain mechanisms of attention – ATTEND (FH, SA, LB), “Fondazione Caritro – Bando Ricerca e Sviluppo Economico” (FH), NIH (DT, MM and KRH: R01 grants EY027314 and EY021209, CVS training grant T32 EY007125), and by an unrestricted grant from the Research to Prevent Blindness (RPB) Foundation to the Flaum Eye Institute. We thank Valeria Piombino for data collection with neurological patients.

## Conflicts of interest

KRH is co-inventor on US Patent No. 7,549,743 and has founder’s equity in Envision Solutions LLC, which licensed this patent from the University of Rochester. The University of Rochester also possesses equity in Envision Solutions LLC. The remaining authors have no competing interests.

